# Investigating a trade-off between the quality of nest grown feathers and pace of development in an altricial bird

**DOI:** 10.1101/2021.04.07.438834

**Authors:** Conor C. Taff, Brianna A. Johnson, Allison T. Anker, Alyssa M. Rodriguez, Jennifer L. Houtz, Jennifer J. Uehling, Maren N. Vitousek

## Abstract

Life history theory provides a framework for understanding how trade-offs generate negative trait associations. Among nestling birds, developmental rate, risk of predation, and lifespan covary, but some associations are only found within species while others are only observed between species. A recent comparative study suggests that allocation trade-offs may be alleviated by disinvestment in ephemeral traits, such as nest-grown feathers, that are quickly replaced. However, direct resource allocation trade-offs cannot be inferred from inter-specific trait-associations without complementary intra-specific studies. Here, we asked whether there is evidence for a within-species allocation trade-off between feather quality and developmental speed in tree swallows (*Tachycineta bicolor*). Consistent with the idea that ephemeral traits are deprioritized, nest-grown feathers had lower barb density than adult feathers. However, despite substantial variation in fledging age among nestlings, there was no evidence for a negative association between developmental pace and feather quality. Furthermore, accounting for differences in resource availability by considering provisioning rate and a nest predation treatment did not reveal a trade-off that was masked by variation in resources. Our results are most consistent with the idea that the inter-specific association between development and feather quality arises from adaptive specialization, rather than from a direct allocation trade-off.

## INTRODUCTION

Understanding the co-expression patterns of ecologically important traits is a primary goal of evolutionary ecology and life history evolution (Stearns 1992; Roff 2002; Agrawal 2020). Theory suggests that trade-offs between different traits should be most pronounced during periods of high demand or low resource availability (van Noordwijk and de Jong 1986). For altricial birds, the early growth period presents a particularly severe challenge, as nestlings must grow quickly to escape the nest and the associated risk of predation while simultaneously developing morphological traits and physiological systems that are critical for lifetime performance (Martin 1995; Martin et al. 2011). Within species, growth rate and time spent in the nest are related to nest predation rates and reduced adult survival, suggesting a resource allocation trade-off (Metcalfe and Monaghan 2003; LaManna and Martin 2016). However, while the relationship between growth and predation risk is also seen in inter-specific comparisons (Bosque and Bosque 1995; Martin 1995; Remeŝ and Martin 2002), the relationship between growth rate and adult survival is not (Martin et al. 2015). Understanding when negative trait associations are similar at different scales (e.g., within-versus between-species) and when those associations occur only at one level, will require a better understanding of the mechanisms that generate these patterns.

Comparative studies of trade-offs have the advantage of being potentially less influenced by the masking effects of between-individual variation in resource availability (van Noordwijk and de Jong 1986; Reznick et al. 2000). However, they also have the disadvantage that the same pattern of inter-specific trait associations can arise by more than one generating mechanism (Figure 1); without complementary intra-specific studies it will often be impossible to distinguish between these possibilities. For example, an apparent trade-off (negative trait association) between species could arise because of a strong constraint resulting in a resource allocation trade-off within each individual of each species, or it could arise due to a “strategic trade-off” resulting from niche specialization or biotic interactions that differ between species. While both of these patterns are often referred to as trade-offs, the mechanism differs (Agrawal 2020). Understanding how and when trait associations translate across scales—from individuals to populations to species—requires both strengthening the scale-dependent predictive framework (reviewed in Agrawal, 2020) and empirical studies that investigate the same trait associations at different scales.

**Figure 1.**
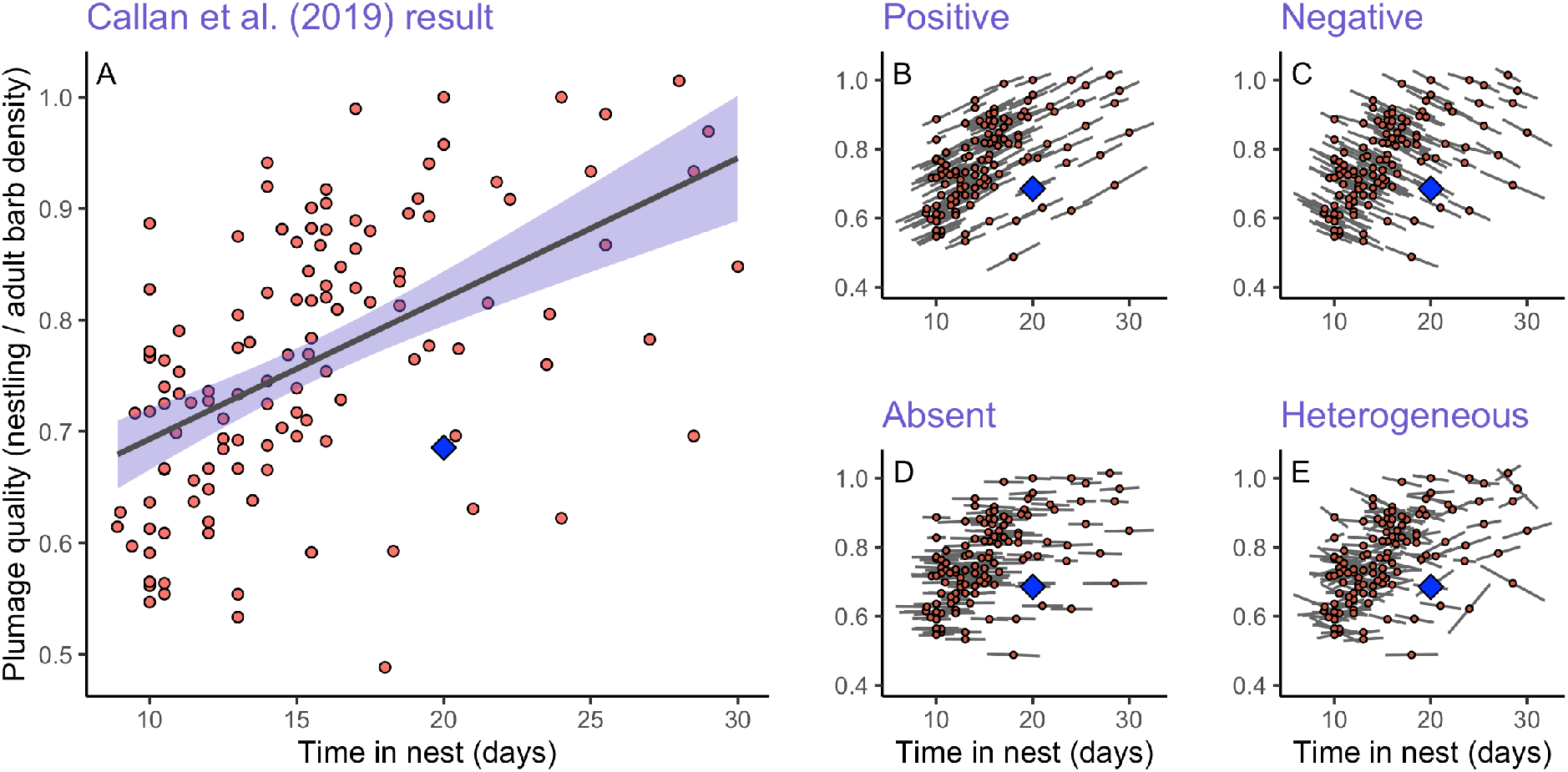
The between-species tradeoff reported in Callan et al 2019 (panel A; redrawn from publicly archived data: Callan et al. 2019*b*) could result from four distinct underlying within species patterns. We used the same mean species points to illustrate hypothetical within species patterns that are positive (B), negative (C), absent (D), or heterogeneous between species (E). These relationships are intended to illustrate the scaling problem when moving from between to within-species inferences and are not based on any biological information about each species shown. In all panels, the value for tree swallows is the large blue diamond.

In the case of nestling growth, Callan et al. (2019) hypothesized that the apparent difference between the growth and adult survival trade-off in comparative versus single species studies might be explained by differences in relative investment in ephemeral traits that contribute relatively little to fitness. They focus in particular on the growth of body feathers in the nest because these feathers are often of lower quality than adult feathers and are replaced shortly after fledging (Rohwer et al. 2005; Butler et al. 2008). Thus, these nest-grown feathers may be relatively unimportant for long term performance and disinvesting in their quality in the nest could ameliorate an otherwise more severe trade-off between growth rate and adult survival. However, Agrawal (2020) argues that negative trait associations often arise between species through adaptive specialization and natural selection even when no direct resource allocation link exists between the two traits; thus inter-specific analysis alone is insufficient to infer within-species trade-offs. In fact, the inter-specific pattern described in Callan et al. (2019) could result from four distinct intra-specific patterns that imply different mechanisms operating at the within-species level (illustrated in Figure 1).

Here, we replicated the comparative analyses of Callan et al. (2019*a*) within a single species—tree swallows (*Tachycineta bicolor*)—to ask whether the correlation between developmental speed and feather quality arises due to a direct resource allocation trade-off. Tree swallow nestlings vary considerably in the exact age of fledging and in the pace of development, even under similar ecological conditions. While nest-grown feathers are quickly molted after fledging, they likely play an important role in thermoregulation for young nestlings, and thermoregulation alone can account for up to a third of all metabolized energy in altricial young (Weathers and Sullivan 1991). Cold snaps are also a major source of mortality in developing tree swallows (Shipley et al. 2020) and feather development in the nest may be particularly important in this species. On the other hand, although predation rates are relatively low in tree swallows, predation is still a common source of mortality, including in our study population (Winkler et al. 2020*a*), and longer development periods lead to an extended period of risk. Thus, it is plausible that a direct resource allocation trade-off could operate on investment in nest-grown feather quality for thermoregulation versus rapid development for predator avoidance.

To test for this trade-off, we measured both feather quality and indicators of developmental rate (mass, wing length, and structural size) in nestling tree swallows. We used a network of automated sensors at each nest to determine the exact age of fledging for each individual nestling. When working at the intraspecific level, resource allocation trade-offs can be masked by variation in resource acquisition (see discussion above; van Noordwijk and de Jong 1986). Thus, we accounted for variation in resource availability in several ways. First, the sensor network provided detailed data on parental provisioning to ask whether the detectability of trade-offs depended on food availability. Second, we took advantage of ongoing experiments in this population that involved experimental predator treatments, which allowed us to assess whether any trade-offs between development and feather quality differed in an environment with high perceived predation risk. Finally, we cross-fostered eggs at each nest before hatching. While cross-fostering was not strictly necessary to test for the trade-off of interest, the ability to flexibly adjust resources during development is a prerequisite of the within-individual trade-off being studied, and cross-fostering allowed us to estimate the extent to which genetic versus environmental effects influenced feather quality.

If the inter-specific association between developmental speed and feather quality results from a resource allocation trade-off that plays out within individuals of each species, then we expected to find an intra-specific association within tree swallows that paralleled the results of Callan et al.’s (2019) comparative study. Specifically, we expected that individual nestlings that fledged at younger ages would pay a cost, as indicated by lower quality nest-grown feathers. Furthermore, we predicted that this trade-off would be most pronounced in nests with either experimentally increased perceived predation pressure or naturally lower parental provisioning rates (*sensu* van Noordwijk & de Jong, 1986). Alternatively, if the inter-specific association arises due to strategic differences in developmental programs as a result of natural selection—rather than a strict resource allocation trade-off—then we would not expect to find a direct trade-off between developmental rate and feather quality within tree swallows.

## METHODS

### General Field Methods

We studied tree swallows breeding near Ithaca, New York, USA in 2018 and 2019 (42.503° N, 76.437° W). During each breeding season (May to July), we monitored every nest at the field sites following established protocols for this long-studied tree swallow population (Winkler et al. 2020*b*). Briefly, each nest box was checked every other day early in the season to determine clutch initiation and clutch completion date to within one day. Around the expected hatching date, we checked boxes every day so that an exact hatch date could be recorded.

Adults at each nest were captured 1-3 times during incubation or provisioning. At the first capture for each adult, we took morphological measurements (mass, flattened wing-chord length, and head + bill length), a blood sample for paternity, and 6-8 feathers from two body regions—the center of the white breast and the rump just above the tail—to measure barb density (see below). For individuals that were not already banded from a previous year, we applied an aluminum USGS band and a passive integrated transponder (PIT) tag that encoded a unique 10-digit hexadecimal string.

Most nests in these two years were part of an experiment that involved targeted manipulations of adult females. While these manipulations were not designed to have any direct effects on nestlings, they may have had indirect effects on nestling resource availability or developmental environment by changing adult provisioning rates, reproductive investment, or antipredator behavior. Therefore, we conducted post-hoc comparisons between nestlings raised in different treatment groups to ask whether adult treatments influenced nestling trade-offs between development and feather quality via changes in resource availability or perceived environmental risk.

The most relevant treatment for the purposes of this study involved 2-3 simulated predation events using a taxidermied mink (*Neovison vision*), a common predator of tree swallows, that occurred shortly before (2018) or shortly after (2019) hatching. In 2018, some females were subject to a challenge treatment that involved a handicapping manipulation where 3 feathers on each wing were bound together with a small plastic zip tie for approximately 5 days late in incubation, thereby reducing flight efficiency and female foraging ability (described in Taff et al. 2019*b*). Finally, females received a social signal manipulation (dulling or sham control of the white breast) in each year (for details of plumage manipulation, see Taff et al. In Press*a*). For both the signal manipulation and challenges, we assumed that any effects on nestlings would primarily occur through altered resources due to provisioning rates, and we therefore analyzed provisioning rates directly from radio-frequency identification (hereafter RFID) records. For the challenge treatments, we also fit models that included the categorical treatments directly as predictors because there is evidence that perceived predation risk can alter developmental trade-offs, independently of its effects on resource availability (e.g., Clinchy et al. 2013; LaManna and Martin 2016).

### Cross-Fostering and Nestling Measurements

One of the goals of this study was to determine the degree to which environmental conditions versus genetic contributions drove differences in feather quality and nestling development. Therefore, we cross fostered eggs from each nest before incubation began so that any contribution to nestling feather development driven by developmental environment (e.g., incubation, provisioning rate) was decoupled from genetic inheritance or maternal effects associated with investment in the egg contents.

We paired nests based on timing of clutch initiation and on day 4 of egg laying we swapped half of the brood between the pair and marked all eggs in each nest with a pencil on the bottom of each egg. At half of the nests, we returned on the following day and swapped the 5^th^ (unmarked) egg between the two nests. This two-step process ensured that some eggs from early and late in the laying order were swapped in case there were differences in yolk contents associated with laying order. In cases where there was not an appropriate nest to swap, we sometimes paired three nests together for cross fostering. A few late season nests did not have any compatible pairs and were not cross fostered.

We banded nestlings when they were 12 days old, collected a blood sample for paternity assignment, and took morphological measurements (mass, wing length, and head + bill length). When nestlings were 15 days old, we once again took a mass measurement, applied a unique PIT tag, and collected feathers exactly as described above for adults to measure barb density. After day 15 we avoided visiting the nest to prevent forced fledging. Final nestling fate and exact fledging date were determined using RFID records and a check of the nest on day 24 to find any nestlings that had died in the nest after day 15.

### RFID Sensor Network

We installed an RFID system at each nest box in the study no later than day 4 of incubation (as in Vitousek et al. 2018). The system consisted of an RFID board held in a waterproof container on the bottom of the nest box (Bridge and Bonter 2011), an antenna that circled the nest box entrance, and a 12-volt battery that powered the system. We programmed the readers to record PIT tags within range (∼2 cm) of the entrance hole every second from 5am to 10pm each day of the breeding season. From raw RFID records, we extracted female and—when possible—male provisioning rates at each nest following the algorithm described in Vitousek et al. (2018).

We also used RFID records to determine the exact age of fledging for each nestling in the population. For each individual nestling, we considered the latest record at the nest box to be the time of fledging. While it is possible that nestlings could leave and then return to the box, we saw no evidence for this behavior in our RFID data even when the sensors were left running long after we had confirmed fledging. Occasionally, RFID units failed because of software problems or dead batteries and we are therefore missing some records from parts of the provisioning period or fledging times for some nestlings.

### Feather Measurements

We measured the density of feather barbs for adults and nestlings following the method developed by Butler et al. (2008) as described in Callan et al. (2019*a*), except that we modified their approach for use with photographs rather than measuring with a dissecting scope. To take photographs, we spread each feather on a microscope slide that had been covered in contrasting cardstock paper with a scale bar. We used black paper as a background when photographing white breast feathers and white paper when photographing brown, green, and blue rump feathers. The feather was pressed down flat with a second clear microscope slide and photographed using a digital camera with a macro lens held in place on a document-scanning platform with diffuse lights. The camera mount ensured that photographs were in sharp focus and always taken at a direct 90° angle from the slide surface to avoid parallax issues when measuring. For each individual, we photographed two breast and two rump feathers.

From the digital photographs, we measured the density of feather barbs using ImageJ (Schneider et al. 2012). We first set the scale for each image using the scale bar that was included in every photograph. Next, we identified the section of the rachis to be measured and marked those points with the annotation tool in ImageJ. For the start point, we chose the most distal point on the feather rachis where a pennaceous barb could be seen branching off from the rachis. For the end point, we chose the most proximal point on the rachis where pennaceous barbs could be clearly seen branching off of the rachis before becoming plumulaceous.

We next measured and recorded the length of the rachis between these two points using the segmented line tool. Finally, we counted the number of pennaceous barbs between the two points and recorded the left and right side barbs separately. We calculated a single barb density measure for the feather by dividing the average count of barbs from the two sides by the length of feather rachis and expressed density in terms of barbs per centimeter of rachis. We repeated this procedure for the two breast and two rump feathers and then averaged the two replicate measurements from each region together to arrive at a single breast density and rump density measurement.

We used multiple feathers from the same bird to estimate biological repeatability across feathers within a subject and multiple measurements of the same photograph by different observers to estimate inter-observer measurement repeatability. In some cases, we did not have complete measurements because we were missing feathers or had only a single feather. Feather measurements were also not available for any nestlings that did not survive to 15 days old.

### Determining Nest of Origin

Adult and nestling blood samples were stored in lysis buffer (Seutin et al. 1991) in the field and DNA was extracted using Qiagen DNeasy Blood & Tissue Kit spin columns following the standard kit protocol. We amplified a set of 9 variable microsatellite loci that have been previously used in this population (Makarewich et al. 2009; Hallinger et al. 2019). Our amplification protocol exactly followed that described in (Hallinger et al., 2019) and details on primer sequences, reaction volumes, cycling conditions, and fragment analysis can be found there. We determined nest of origin by comparing nestlings to their putative mothers (the females from the 2-3 nests in each cross-fostering pair). Nestlings that matched only one putative mother at 8 of 9 loci were considered to have been laid by that female. Using these criteria, we were able to assign definitive genetic mothers to 275 of 313 sampled nestlings. The remaining 38 nestlings were either missing blood samples, missing maternal genotype information, mismatched at more than two loci, or had two putative mothers that were so similar that the genetic mother could not be determined definitively. Those nestlings are excluded from all analyses that included nest of origin but included in summary statistics.

### Data Analysis

We assessed repeatability between observers and between multiple feathers from the same individual by estimating the intra-class correlation coefficient (ICC) in linear mixed models with no covariates as implemented by the ‘rptR’ package version 0.9.22 in R (Stoffel et al. 2017). Overall differences in adult and nestling feather barb density measurements were assessed using linear mixed models with age (adult or nestling) as a fixed effect and nest as a random effect. To assess the extent to which environment (nest identity) and genetics (genetic mother) jointly contributed to variation in nestling feather barb density, we fit LMMs using the ‘rptR’ package that included random effects for both the nest identity and the genetic mother. We report the ICC for each of these effects as an estimate of the amount of variation explained by nest and genetic mother.

To evaluate the evidence for a trade-off between the speed of development and feather quality, we fit a series of models that included proxies of developmental pace as response variables (exact age at fledging, mass and morphological measurements on day 12 or 15). For fixed effects, these models included breast and back feather density, a categorical effect for the challenge treatment that the female received (control, handicap, or predator), and two-way interactions between feather density measurements and challenge treatment. For simplicity, we present reduced models that exclude interactions receiving no support. These models also included nest and genetic mother as random effects to account for the non-independence of nestlings from the same nest or mother.

Finally, we fit models to ask whether variation in female provisioning rate predicted nestling back or breast feather density (male provisioning was not considered because not all males had PIT tags). We first estimated standardized female provisioning rate by fitting a model of daily provisioning that included brood size, nestling age, a quadratic effect of nestling age, and random effects for the day of year (to account for weather differences) and nest identity (to account for repeated observations at each nest). We extracted a provisioning rate for each female from this model and standardized to a mean of 0 and standard deviation of 1. This standardized provisioning rate was then used as a predictor of nestling feather barb density in an LMM with nest identity and genetic mother included as random effects as described above.

All LMMs apart from those used to calculate ICC were fit with the ‘lme4’ package version 1.1-26 in R (Bates et al. 2015). Predictors were considered to be meaningful if the confidence interval did not cross zero. In tables presenting mixed model details, we also include p-values based on the Satterthwaite approximation as implemented by ‘lmerTest’ version 3.1-3 (Kuznetsova et al. 2017). All figures and analyses were produced in R version 4.0.2 (R Core Development Team 2020).

## RESULTS

In total, our analysis included 313 nestlings raised in 85 nests with at least one feather region measurement and 274 adults with at least one feather region measurement. In a validation dataset, inter-observer repeatability of barb density measurements from the same feather photograph was high (*n* = 149 measurements of 38 photographs by 8 observers; repeatability = 0.96; 95% CI = 0.94 to 0.98). Measurements of two independent feathers from the back or the breast of the same bird were also repeatable (breast: *n* = 1123 measures of 572 individuals; repeatability = 0.78; CI = 0.74 to 0.81; back: *n* = 1021 measures of 572 individuals; repeatability = 0.68; CI = 0.63 to 0.72).

Overall, there was a moderate positive correlation between barb density of back and breast feathers within an individual (Figure 2). This relationship was observed for both adults and nestlings with a similar slope in each group (Pearson’s correlation for adults and nestlings combined: *r* = 0.66, CI = 0.61 to 0.70; adults *r* = 0.34, CI = 0.23 to 0.45; nestlings *r* = 0.24, CI = 0.13 to 0.35).

**Figure 2.**
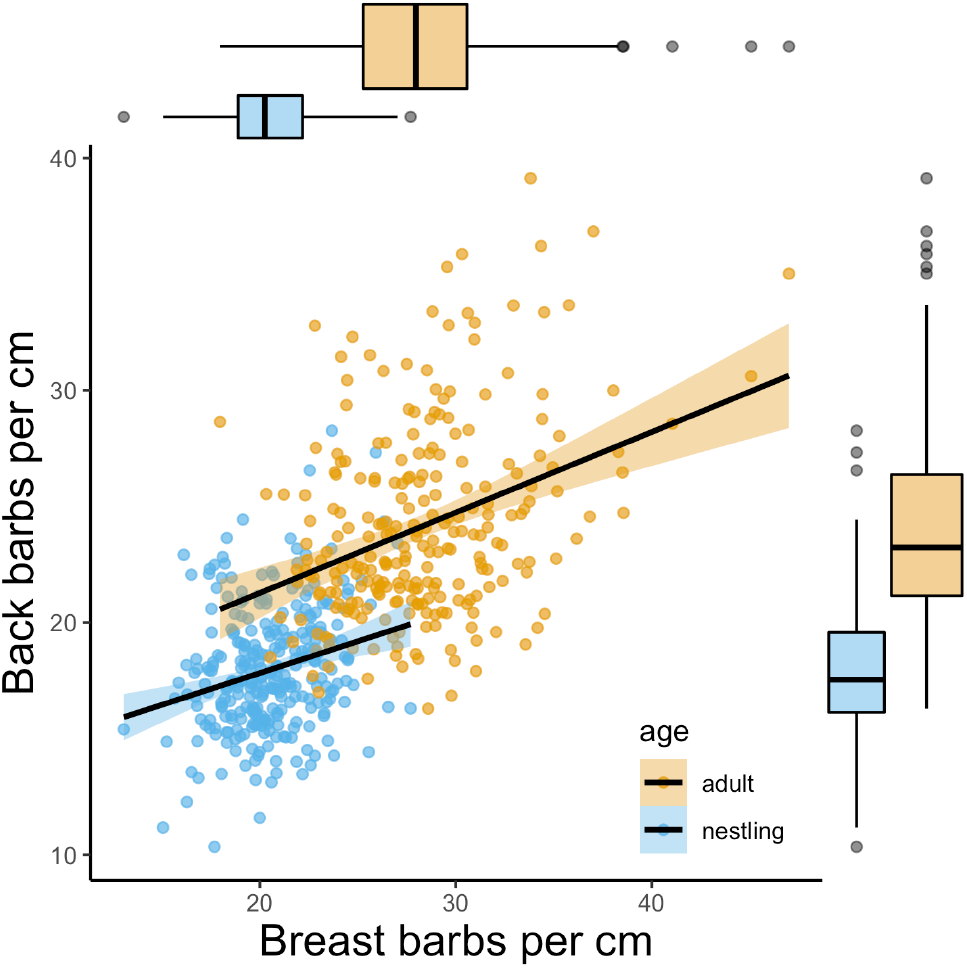
Relationship between breast and back barb density for feathers measured from the same individual for adults (orange) and nestlings (blue). Box and whisker plots in the margins show the distribution of barb density measurements for each body region and age group.

As expected, adults had substantially higher barb density for both back and breast feathers (Figure 2). For back feathers, nestlings had an overall barb density of 17.9 barbs per cm, while adults had 24.3 barbs per cm (LMM with nest as a random effect, ® for nestlings = −6.39, CI = −6.97 to −5.81). For breast feathers, nestlings had an overall barb density of 20.4 barbs per cm, while adults had 28.2 barbs per cm (LMM ® for nestlings = −7.79, CI = −8.33 to −7.26). Thus, nestlings had on average 73.7 % (back) and 72.4 % (breast) of the barbs per cm as adults.

### Environmental and Genetic Influence on Barb Density

For breast feathers, variation in nestling barb density was explained by both the genetic mother and the nest environment that a nestling was raised in. The adjusted ICC of nest environment controlling for genetic mother in an LMM was 0.33 (CI = 0.2 to 0.442). The adjusted ICC of genetic mother controlling for nest environment in an LMM was 0.27 (CI = 0.14 to 0.39). For back feathers, genetic mother explained some variation in nestling barb density, but nest environment explained little. The adjusted ICC of genetic mother for back feather barb measurements controlling for nest environment was 0.20 (CI = 0.05 to 0.34). The adjusted ICC of nest environment controlling for genetic mother was 0.07 (CI = 0.0 to 0.19).

For breast measurements, unadjusted ICC estimates were nearly identical, but for back measurements, unadjusted estimates were higher for both categories, suggesting that nest environment and genetic mother explained much of the same variation. Unadjusted ICC for genetic mother on back barbs was 0.26 (CI = 0.12 to 0.39) and for nest environment was 0.22 (CI = 0.08 to 0.34).

### Barb Density by Time in the Nest

Nestlings fledged an average of 22.6 days after hatching (standard deviation = 1.5, range = 18 to 29 days). There was no apparent relationship between age at fledging and the density of feather barbs for either back or breast feathers (Figure 3). There was also no indication that the (lack of) relationship differed by treatment group (Table 1; unsupported interactions are not shown). However, there was a main effect of the predator treatment with nestlings from the predation treatment fledging at older ages than those from control or handicap nests. Nest identity explained much of the variation in fledging age and genetic mother explained little additional variation (Table 1; marginal R^2^ for reduced model = 0.095, conditional R^2^ including random effects = 0.657).

**Figure 3.**
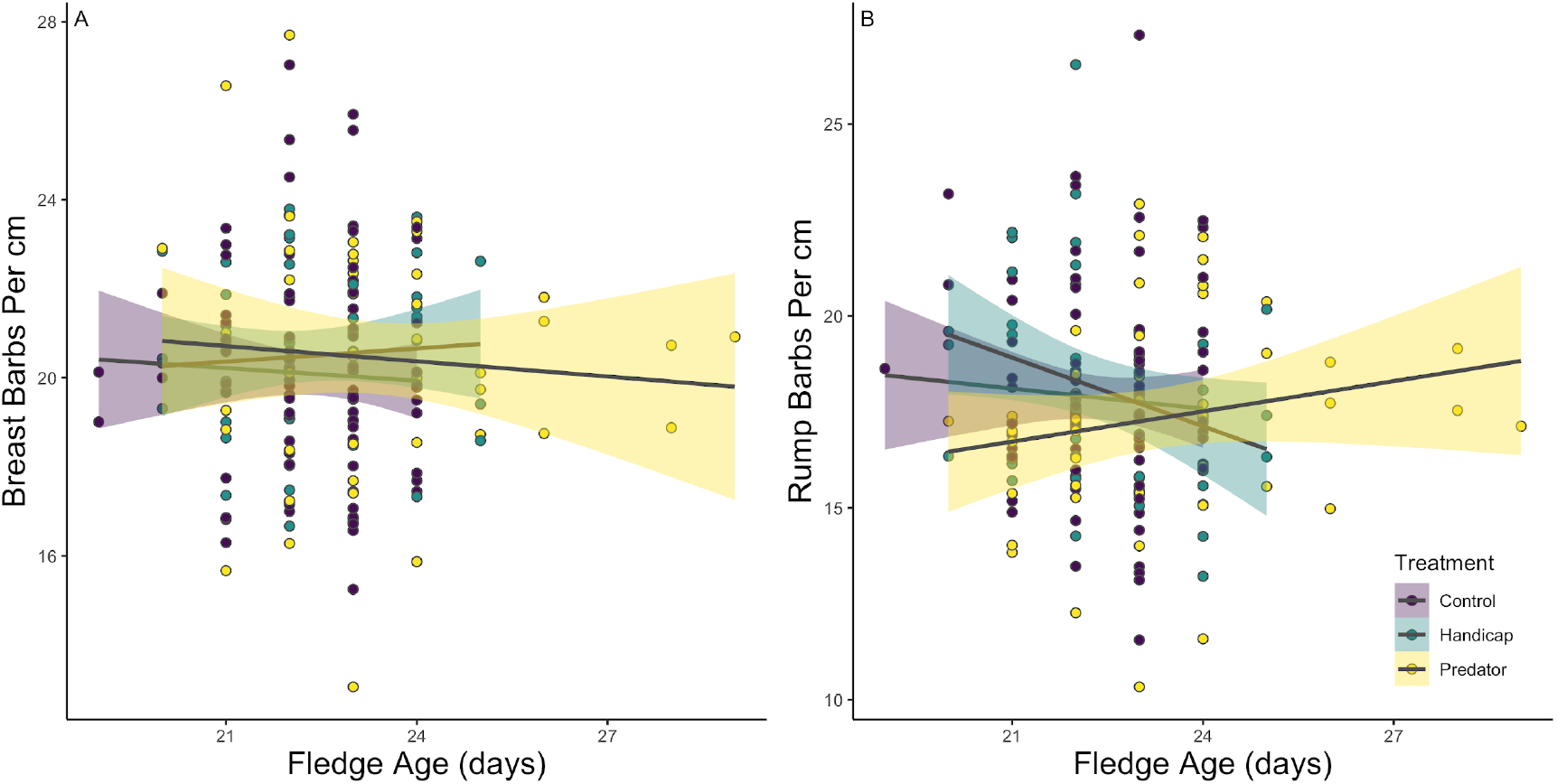
Fledging age was not related to breast (A) or back (B) feather barb density per cm for nestlings in the three different treatment groups.

**Table 1.** Feather barb density and nest treatment as predictors of fledging age.

| *Predictors* | **Fledge Age (Days)** |  |  |
| --- | --- | --- | --- |
|  | *Estimates* | *CI* | *p* |
| Intercept (Control) | 22.40 | 20.35 – 24.45 | **<0.001** |
| Back Barb Density | 0.04 | -0.04 – 0.11 | 0.328 |
| Breast Barb Density | -0.04 | -0.13 – 0.05 | 0.410 |
| Handicap | -0.06 | -0.99 – 0.88 | 0.905 |
| Predator | 1.17 | 0.33 – 2.02 | **0.006** |
| **Random Effects** |  |  |  |
| $\sigma^2$ | 1.00 | | |
| $\tau_{00}$ Mother | 0.01 | | |
| $\tau_{00}$ Nest | 1.63 | | |
| ICC | 0.62 |  |  |
| N<sub>Nest</sub> | 60 |  |  |
| N<sub>Mother</sub> | 75 |  |  |
| Observations | 177 |  |  |
| Marginal R<sup>2</sup> / Conditional R<sup>2</sup> | 0.095 / 0.657 |  |  |

### Correlation Between Nestling Feather Barb Density and Morphology

Breast feather barb density was positively related to day 12 and day 15 nestling mass, but only in the predator treatment group (Figure 4, Table 2). Back barb density was not related to nestling wing length, head plus bill length, or mass in any treatment group. Overall, nestlings raised in the predator treatment group had shorter wings on day 12 and lower mass on day 12 and 15 than did nestlings raised in either the handicap or control group (Table 2). However, the amount of variation in mass explained by feather measurements was small compared to that explained by random effects fitted for the nest environment and the genetic mother (Table 2; marginal R^2^ of main effects = 0.02 to 0.09; conditional R^2^ including random effects = 0.32 to 0.70).

**Figure 4.**
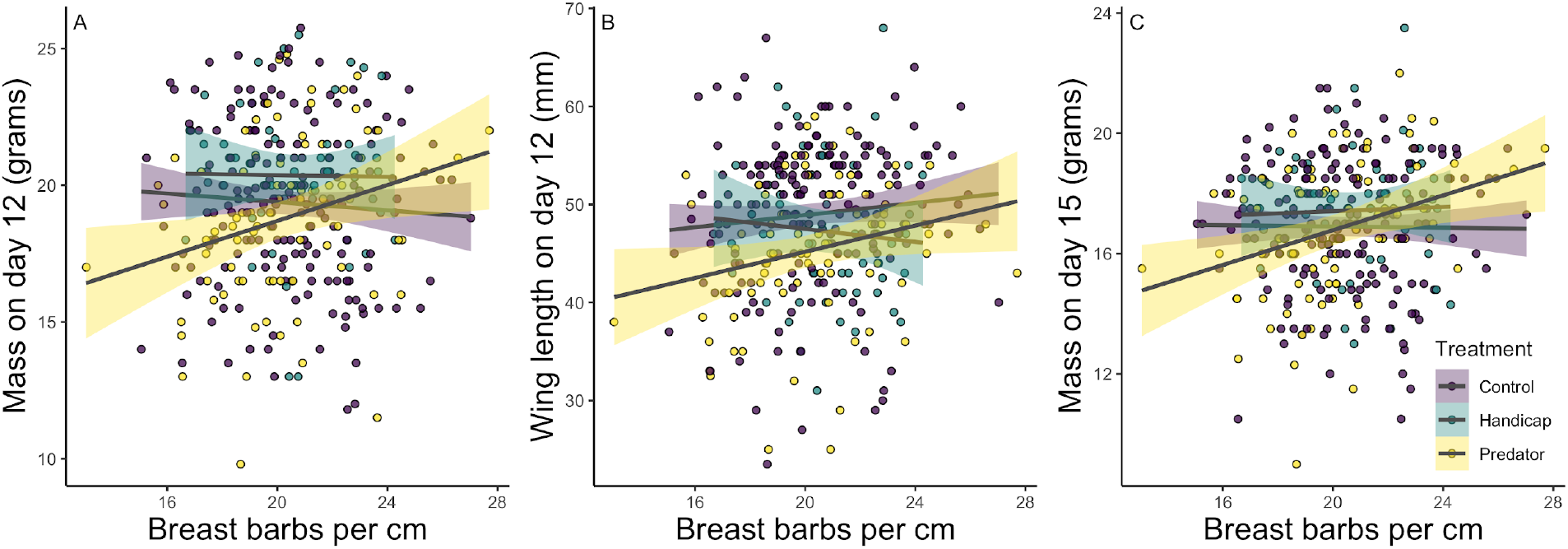
Nestling mass on day 12 (A), flattened wing chord length on day 12 (B), and mass on day 15 (C) in relation to breast feather barb density by treatment group.

**Table 2.** Linear mixed models showing relationship between barb density and nestling morphology by treatment group. Genetic mother and nest of development are included as random effects.

| *Predictors* | **Day 12 Mass** |  |  | **Day 12 Head + Bill** |  |  | **Day 12 Flat Wing** |  |  | **Day 15 Mass** |  |  |
| --- | --- | --- | --- | --- | --- | --- | --- | --- | --- | --- | --- | --- |
|  | *Estimates* | *CI* | *p* | *Estimates* | *CI* | *p* | *Estimates* | *CI* | *p* | *Estimates* | *CI* | *p* |
| Intercept | 21.92 | 18.26 – 25.57 | **<0.001** | 24.20 | 22.80 – 25.60 | **<0.001** | 48.71 | 39.91 – 57.51 | **<0.001** | 17.75 | 14.76 – 20.74 | **<0.001** |
| Back Barb Density | -0.04 | -0.14 – 0.07 | 0.495 | 0.01 | -0.04 – 0.06 | 0.813 | 0.05 | -0.20 – 0.30 | 0.677 | -0.02 | -0.11 – 0.07 | 0.649 |
| Breast Barb Density | -0.10 | -0.26 – 0.06 | 0.221 | 0.03 | -0.03 – 0.09 | 0.281 | -0.03 | -0.42 – 0.35 | 0.864 | -0.03 | -0.16 – 0.11 | 0.711 |
| Handicap | 3.43 | -4.55 – 11.41 | 0.399 | -0.03 | -0.55 – 0.49 | 0.913 | 7.73 | -11.30 – 26.75 | 0.426 | -0.69 | -7.24 – 5.87 | 0.838 |
| Predator | -6.27 | -12.15 – -0.39 | **0.037** | -0.34 | -0.77 – 0.10 | 0.132 | -15.83 | -30.01 – -1.64 | **0.029** | -4.72 | -9.38 – -0.06 | **0.047** |
| Breast Barb * Handicap | -0.11 | -0.48 – 0.27 | 0.578 |  |  |  | -0.40 | -1.29 – 0.49 | 0.376 | 0.07 | -0.24 – 0.38 | 0.674 |
| Breast Barb * Predator | 0.28 | 0.00 – 0.56 | **0.049** |  |  |  | 0.53 | -0.14 – 1.21 | 0.122 | 0.23 | 0.01 – 0.46 | **0.041** |
| **Random Effects** |  |  |  |  |  |  |  |  |  |  |  |  |
| $\sigma^2$ | 3.35 | | | 0.99 | | | 18.47 | | | 2.50 | | |
| $\tau_{00}$ | 1.33 | Mother | | 0.02 | Mother | | 5.14 | Mother | | 0.72 | Mother | |
|  | 3.61 | Nest |  | 0.42 | Nest |  | 33.02 | Nest |  | 1.33 | Nest |  |
| ICC | 0.60 |  |  | 0.31 |  |  | 0.67 |  |  | 0.45 |  |  |
| N | 84 | Nest |  | 84 | Nest |  | 84 | Nest |  | 82 | Nest |  |
|  | 90 | Mother |  | 90 | Mother |  | 90 | Mother |  | 89 | Mother |  |
| Observations | 290 |  |  | 290 |  |  | 288 |  |  | 286 |  |  |
| Marginal R<sup>2</sup> / Conditional R<sup>2</sup> | 0.048 / 0.615 |  |  | 0.020 / 0.320 |  |  | 0.085 / 0.702 |  |  | 0.031 / 0.467 |  |  |

After controlling for nestling age and brood size, female provisioning rate was not associated with nestling back feather barb density (LMM with nest and genetic mother as random effects; full model marginal R^2^ = 0.008; effect of standardized provisioning rate = −0.26; CI = −0.66 to 0.15). However, a higher rate of female provisioning was associated with decreased nestling breast feather barb density (Figure 5, LMM full model marginal R^2^ = 0.04; effect of provisioning rate = −0.49; CI = −0.87 to - 0.10).

**Figure 5.**
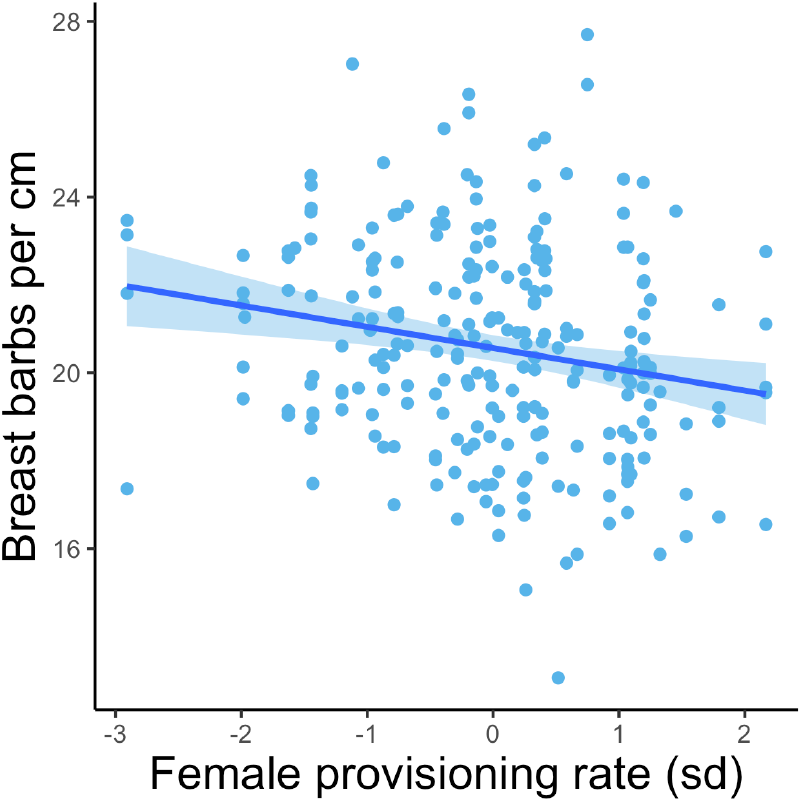
Nestling breast barbs per centimeter in relation to standardized female provisioning rate. Provisioning rate accounts for nestling age and brood size (see methods) and is shown in units of standard deviations.

## DISCUSSION

Consistent with the idea that ephemeral traits may be deprioritized during development, we found that tree swallow nestlings grew body feathers that had lower barb density than those of adults from the same body regions. Individuals varied considerably in their barb density, but there was high repeatability within individuals for feathers from the same body region and a moderate association between feathers from different body regions. Cross fostering revealed that there was both a strong environmental and genetic basis to variation in feather barb density. Despite the fact that there was substantial variation in both feather quality and in the pace of development—measured as either the exact age of fledging or body size—we found no evidence for a within-individual resource allocation trade-off between feather quality and rapid development. Moreover, accounting directly for variation in resource acquisition by including maternal provisioning rates or experimental challenges on parents did not reveal any hidden trade-off, as might be expected if variation in resource acquisition had masked allocation trade-offs (sensu van Noordwijk & de Jong, 1986). In fact, higher female provisioning rates were associated with slightly lower feather quality and, among nestlings in a nest predation treatment group, there was a positive association between feather quality and body size measurements; these patterns are opposite to those predicted if reduced resources had forced differential investment in feathers versus growth. Our results highlight the difficulty of inferring individual level resource trade-offs from comparative data alone.

In a comparative study of feather barb density from nest-grown feathers including 123 species, Callan et al. (2019) found a strong relationship between time in the nest and feather quality (Figure 1A). Species that develop more rapidly in the nest have consistently lower quality feathers—as compared to adults of the same species— than do those that develop more slowly in the nest. The authors hypothesize that this relationship arises from a resource allocation trade-off, with species that must develop rapidly in order to leave the nest and avoid predation shunting resources away from other traits. They argue that these trade-offs may be especially apparent in ephemeral traits, like nest-grown feathers, because the fitness consequences of lower quality ephemeral traits are relatively low since they are quickly replaced after fledging. How can we resolve this evidence for an apparently strong trade-off between development and feather quality among species with the lack of evidence for a trade-off found within a single species in our study? Agrawal argues that often, “trade-offs at one level of organization will provide little insight into what may occur at other levels” (pg. 3; Agrawal, 2020) without additional consideration of the mechanism that generates trait associations. His review goes on to develop a conceptual framework for considering what types of trait associations (or trade-offs) are likely to scale across levels or to only be apparent at a single level.

Interpreted in light of this framework, our results in tree swallows and the comparative results presented by Callan et al. (2019) are most consistent with the idea that the inter-specific association between rapid development and feather quality arises from adaptative specialization to different developmental strategies by each species, rather than from a direct resource allocation trade-off that plays out within species or individuals (Figure 1D). This is perhaps unsurprising given the nature of the proposed trade-off. Direct resource allocation trade-offs are most likely to arise with strong constraints and between life history traits that are directly associated with fitness when all else is held equal (Agrawal, 2020; Roff, 2002; Stearns, 1992). In contrast, negative trait associations among species often represent the outcome of adaptive specialization based on strategic optimization (Futuyma and Moreno 1988; Agrawal 2020). In the case of ephemeral feathers, the direct developmental costs of feather barbs may not be severe enough to generate a within species resource allocation trade-off and may not share a direct mechanistic link with the overall rate of development (e.g., a shared developmental pathway). Additionally, plastic responses are only favored when relevant information is available (Pigliucci 2005), and if nestlings cannot reliably assess their own developmental pace, there may be no realized benefit to modifying the allocation of resources to faster development over feathers. In contrast, when comparing different species, strong selection for particular developmental trajectories (e.g., fast fledging to avoid predation) may result in subsequent selection to optimize investment in secondary traits, such as ephemeral feather quality, without the need to invoke a direct allocation trade-off.

Alternatively, it is possible that some species do face direct resource allocation trade-offs between feather quality and developmental rate, but that tree swallows are not representative of a more general mechanism (Figure 1E). Several aspects of tree swallow life history might make them less likely to show this particular allocation trade-off. First, as cavity nesters, tree swallows experience relatively low predation rates and spend an unusually long time in the nest for their size (Winkler et al. 2020*b*), which may mitigate the need to redirect resources towards rapid development. Second, cold snaps create strong selection events for tree swallow nestlings because the parents are entirely dependent on flying insects (Winkler et al. 2013). When flying insects are scarce, parents travel far from the nest and are absent for long periods (Stocek 1986). These cold snaps can lead to mass mortality events for nestlings, but after feathers are grown nestlings are much less vulnerable because they can thermoregulate independently (Shipley et al., 2020). Thus, for tree swallows in particular, growing effective feathers in the nest might be especially important even when resources are scarce. It is possible that direct resource allocation trade-offs might be observed in other species that face different life history challenges.

We also found that feather barb density itself and morphological relationships with barb density differed for breast versus rump feathers. In general, breast feathers had higher barb density and were more clearly influenced by both nest environment and genetic mother; feathers from this body region were also the only ones that showed any relationship with nestling morphological measures. In adult tree swallows, feathers in both of these body regions are putative social signals, but their function and color patterns differ dramatically. Breast feathers are light gray to pure white and have been implicated in social signaling and aggression (Beck et al. 2015; Taff et al. 2019*b*). Rump feathers are brown to iridescent blue-green and have been associated with mate choice and extrapair paternity in males (Bitton et al. 2007; Van Wijk et al. 2016; Whittingham and Dunn 2016). Adult female rump feathers go through a delayed plumage maturation with one year old females displaying brown feathers that turn iridescent blue-green in subsequent years and this process may also mediate social relationships and performance (Berzins and Dawson 2016; Dakin et al. 2016). We do not know at present whether the color of these patches has a function in fledgling tree swallows and, if so, how that might relate to barb density. From a thermoregulation perspective, the importance of breast and rump feathers may also differ. For nestlings huddling in a nest cup, the rump feathers are more directly exposed to ambient air, but the breast feathers insulate the pectoralis muscles, which is a major source of shivering thermogenesis in birds (West 1965). We also observed substantial variation in the age of feathering in between nestlings in our study; some nestlings had fully developed feathers across most of the body by day 15 while others had only small pin feathers. Because feathers are inert after exiting the follicle, the barb measurements we took from 15-day-old nestlings represent the influence of conditions experienced earlier in development. It seems likely that overall feathering is more important for thermoregulation than barb density *per se*, but we do not know at present whether the quality of individual feathers and overall rate of feather growth are mechanistically linked.

Although we did not find any evidence for an allocation trade-off, we did find that predator treatments resulted in an association between feather barb measurements and body size in some cases. In many species, experimentally increasing the perceived threat of predation leads to faster nestling development (LaManna and Martin 2016). In contrast, nestlings in our study fledged later when raised in a nest that experienced predation treatments. While we cannot be certain about the cause of this discrepancy, it seems likely that the details of our treatment may have contributed to the difference. Because our predation treatments were targeted at adult females and occurred shortly before or after hatching, there was little or no opportunity for nestlings to directly perceive predator treatments. Rather, any perceived threat would have occurred through the indirect effect of subsequent parental behavior. In contrast, previous studies targeting nestlings have manipulated cues (e.g., auditory or visual presentations) that nestlings could perceive directly (Hallinger et al., 2019; LaManna & Martin, 2016). In our study, nestlings in the predation treatment group showed a positive correlation between body size and breast feather density (the opposite to that predicted if predation threat increased relative investment in growth or decreased acquisition of resources). There are two possible explanations for this pattern. First, parental behavior may have been altered in a way that impacted nestling developmental trajectories. While we did not observe differences in overall provisioning rate, parents may have changed brooding schedules, or the type or quality of food delivered to nestlings. Second, as a result of altered parental behavior, the feather relationships we observed in the predation group may be the result of differential survival to day 15 rather than changes in relative resource allocation. Because we only had feather measurements for nestlings that lived to day 15, we cannot assess whether differential survival contributed to the patterns that we observed. In either case, our results do not support a role for predation threat in revealing a hidden resource allocation trade-off between development and feather quality in tree swallows, although variation in predation risk is clearly an important driver of trait correlations in inter-specific comparisons (Callan et al. 2019*a*).

Our results highlight the fact that strong trait-correlations among species do not necessarily scale across levels of biological organization and that it is often difficult to infer within-species mechanisms from among species associations. Moving forward, we suggest that more studies are needed that integrate measures of trait covariation within individuals and species with comparative analyses in order to parse the hierarchical nature of variation in trait correlations. These approaches are complementary: within-species measurements are needed to identify the proximate mechanisms producing trait correlations, while comparative studies are needed to understand how evolution has shaped population and species level strategies and to understand the conditions under which trade-offs represent strong constraints on life history evolution. Combining these approaches is a critical step towards developing a predictive framework for understanding when and why trait-associations do or do not scale across levels of biological organization.

## ETHICAL NOTE

We received approval for all of the procedures described here from the Cornell University Institutional Animal Care & Use Board (IACUC protocols # 2001-0051 and 2019-0023). Sampling and capture in the field were approved by federal and state permits.

## ACKNOWLEDGMENTS

We thank the field and lab technicians who helped to collect data for this project, including Bashir Ali, Paige Becker, Raquel Castromonte, KaiXin Chen, Jeremy Collison, Alex Dopkin, Zapporah Ellis, Audrey Fox, Sungmin Ko, Raisa Kochmaruk, Christine Kallenberg, Brittany Laslow, Alex Lee-Papastavros, Jabril Mohammed, Yusol Park, Callum Poulin, Emma Regnier, Bella Somoza, and Kwame Tannis. We also thank Anurag Agrawal and Vanya Rohwer for comments on early versions of the manuscript.

Research funding was provided by NSF-IOS 1457251, a USDA Hatch Grant, and DARPA YFA D17AP00033 (to MNV). Support for BAJ, ATA, and AMR was provided by the Cornell Lab of Ornithology and NSF-REU funds. JLH and JJU were supported by NSF GRFP DGE-1650441.

## DATA AND CODE ACCESSIBILITY

The complete dataset and code required to reproduce all analyses and figures is available on GitHub (https://github.com/cct663/tres_feather_density).

## AUTHOR CONTRIBUTION STATEMENT

ATA, BAJ, CCT, and MNV contributed to the conceptualization of the study. AMR, ATA, BAJ, CCT, JJU, and JLH contributed to data collection. AMR, ATA, and BAJ, carried out measurements for the study. MNV contributed funding acquisition. CCT conducted analysis and visualization with input from all authors. CCT wrote the original draft of the manuscript with BAJ and with subsequent feedback and editing from all authors. All authors approved the final version of the manuscript.

